# Methionine metabolism influences the genomic architecture of H3K4me3 with the link to gene expression encoded in peak width

**DOI:** 10.1101/243196

**Authors:** Ziwei Dai, Samantha J. Mentch, Xia Gao, Sailendra N. Nichenametla, Jason W. Locasale

## Abstract

Nutrition and metabolism are known to influence chromatin biology and epigenetics by modifying the levels of post-translational modifications on histones, yet how changes in nutrient availability influence specific aspects of genomic architecture and connect to gene expression is unknown. To investigate this question we considered, as a model, the metabolically-driven dynamics of H3K4me3, a histone methylation mark that is known to encode information about active transcription, cell identity, and tumor suppression. We analyzed the genome-wide changes in H3K4me3 and gene expression in response to alterations in methionine availability under conditions that are known to affect the global levels of histone methylation in both normal rodent physiology and in human cancer cells. Surprisingly, we found that the location of H3K4me3 peaks at specific genomic loci was largely preserved under conditions of methionine restriction. However, upon examining different geometrical features of peak shape, it was found that the response of H3K4me3 peak width encoded almost all aspects of H3K4me3 biology including changes in expression levels, and the presence of cell identity and cancer associated genes. These findings reveal simple yet new and profound principles for how nutrient availability modulates specific aspects of chromatin dynamics to mediate key biological features.

## Introduction

Genes interact with environmental factors such as nutrition to shape the epigenome that together influences gene activity and organismal physiology. Metabolism is also shaped by genes and environment and has a substantial contribution to epigenetics^1–5^. This nexus is essential in numerous biological contexts including maintaining different stages of pluripotency^6–14^, mediating an immune response^15–18^, promoting or suppressing cancer progression^19–25^, and transducing information about metabolic health and longevity from parent to offspring^26–29^. The molecular foundation of this interaction is in large part determined by the modifications on chromatin.

Chromatin is affected by metabolism through changes in the concentrations of metabolites that serve as substrates and cofactors for post-translational modifications. These concentrations are dynamic and are mediated by changes in metabolic pathway activity or flux that arise from transcriptional programs and nutrient availability. For example, histone methylation requires S-adenosylmethionine (SAM) as the universal methyl donor. SAM is derived from methionine^30^ and its concentration can fluctuate in physiological conditions around values that can limit the activity of histone methyltransferases^31^.

In plasma, methionine is in some reports the most dynamic of the 20 amino acids and the variation can to a large extent be explained by diet^32^. Recently work from us and others has shown that dietary modulation of methionine concentrations that approach the lower end of what can be observed in humans leads to bulk changes in the levels of histone methylation^6,32^. Other studies have reported similar findings in that changes to SAM levels or to the levels of alpha-ketoglutarate that modify the activity of demethylase enzymes induce global changes in the levels of histone modifications^7,10,12,15,22,33–38^. When these modifications are known to mark key aspects of chromatin status, global changes could have broad consequences to epigenomic programs. How these bulk changes to the levels of post-translational modifications on chromatin alter the genomic architecture of histone marks and relate to gene expression is, however, largely unknown.

One attractive model to investigate this interaction at the genome scale is the tri-methylation of Histone H3 on lysine 4 (H3K4me3). The global (i.e. bulk) levels of this mark are dynamically and reversibly responsive to the levels of methionine^32^. In addition, there are numerous lines of evidence indicating that the structural features of H3K4me3, such as the width or breadth of the peak as deposited over a genic region, encode information such as gene activity, and gene function such as the presence of a developmental program, cell type identity, or a tumor suppressor^39–43^. Thus, changes in H3K4me3 may be relevant to developmental transitions and tumor suppression. How metabolic dynamics that occur due to differences in nutritional status or metabolic pathway activity might affect these programs and gene activity related to H3K4me3 is largely unknown.

We have shown previously that methionine availability modulates bulk levels of H3K4me3 by modifying SAM concentrations^32^. In this present study, we questioned whether changes in methionine availability that are known to affect global levels of H3K4me3 affect specific aspects of the genomic architecture and gene expression regulation. We considered a mouse model of dietary methionine restriction leading to changes in bulk levels of H3K4me3 in liver and cultured human cancer cells (HCT116) subjected to methionine restriction in culture media that also leads to changes in bulk levels of H3K4me3. We studied genome-wide H3K4me3 dynamics using a quantitative ChIP-seq analysis that considers peak geometry and characterized the connection to gene expression dynamics. We found that aspects of the peak geometry such as the height and area were overall reduced that accounted for most of the global changes. Strikingly, however, while the most conserved feature of H3K4me3 dynamics was found in the peak width, changes in peak width but not other features of peak geometry were associated with nearly all aspects of H3K4me3 biology including cell identity related gene expression programs and the dynamics of gene expression.

## Results

### Methionine restriction reduces global levels of H3K4me3 but maintains its genomic distribution

To begin to study the impact of methionine availability on the genomic architecture of H3K4me3, we applied chromatin immunoprecipitation with sequencing (ChIP-seq) to map genomic locations enriched with H3K4me3 in HCT116 cells cultured under high (100 μM) and low, methionine-restricted (MR) (3 μM) methionine availability which is known to lead to several-fold global changes in the bulk levels of H3K4me3^32^. Aligning the reads to a reference genome followed by peak calling identified a set of H3K4me3 peaks, i.e., genomic regions significantly enriched (MACS2 enrichment Q-value<1e-5) with the ChIP-seq reads in comparison with the control (Fig 1A). Comparing total peak number (coefficient of variation (CV) = 0.01, Fig 1B), genomic location (Jaccard index = 0.88, Fig 1C) and the set of genes marked by peaks (Jaccard index = 0.92, Fig S1A) between high and low methionine conditions showed that the distribution of H3K4me3 genomic locations was highly conserved in response to MR (Fig 1B, 1C) while an overall reduction in H3K4me3 in peak intensity also observed (Fig S1B, S1C). Genes with gained or lost H3K4me3 peaks in response to MR tended to be observed in smaller peaks (Wilcoxon rank-sum P-value < 1e-4 for 5 out of the 6 comparisons, Fig S1D, S1E) and were not significantly enriched with specific biological functions compared to randomly chosen gene sets (median(-log_10_(enrichment Q-value)) = 4.25 for gained peaks compared to 7.76, 8.05 and 8.98 for three random gene sets of identical size, Fig S1F, and median(-log_10_(enrichment Q-value)) = 3.00 for lost peaks compared to 7.12, 6.05 and 5.08 for three random gene sets with identical size, Fig S1G) and thus likely attributable to technical noise. We also analyzed the composition of genomic elements covered by the peaks and found, as has been reported with H3K4me3^44^, that most peaks (11210 peaks, 81%) covered promoter regions with a smaller subset of peaks (2599 peaks, 19%) that appeared on non-promoter regions such as intergenic regions and introns (Fig S1H). To further investigate these changes, we next developed several quantitative descriptors for individual peaks (Fig 1D). We used height, area and width to evaluate the size of the peaks and compared these metrics in high and low methionine conditions. Genomic regions in the high and low methionine conditions were merged (Fig 1A) to include less abundant peaks called only in one condition in the analysis. All three peak size descriptors showed high correlations (Spearman’s rank correlation coefficient>0.95 and random permutation test P-value<1e-323) between high and low methionine conditions (Fig 1E), implying that the overall H3K4me3 landscape (i.e. relative size of each individual peak) is robust to methionine availability (linear regression coefficient = 0.87 for width, 0.85 for height and 0.70 for area).

**Figure 1.**
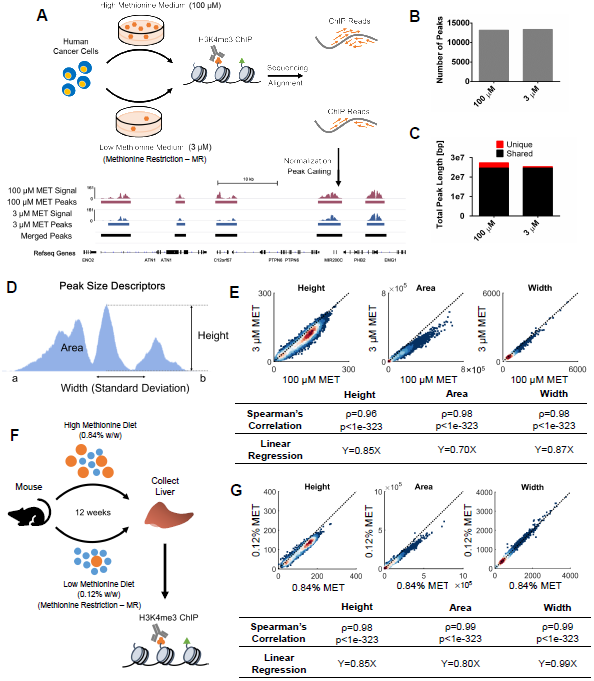
Methionine restriction reduces global levels of H3K4me3 but maintains its genomic distribution. A. MR experiment and ChIP-seq data analysis pipeline in human cancer cells HCT116. B. Number of H3K4me3 peaks called under high (100 μM) and low (3 μM) methionine conditions in human cancer cells. C. Total length of shared and unique peak regions for high and low methionine conditions in human cancer cells. D. Definition of peak height, area and width. E. Density scatter plots comparing H3K4me3 peak heights, areas and widths between high and low methionine conditions in human cancer cells. Colors represent for dot density. Dashed lines show identical x and y coordinates. Spearman’s rank correlation coefficients between the high and low methionine conditions and linear regression coefficients are shown in the table at bottom. F. MR experiments scheme *in vivo*. G. Density scatter plots comparing H3K4me3 peak heights, areas and widths in mouse liver between high (0.84% w/w) and low (0.12% w/w) methionine diets.

To determine whether this conservation in H3K4me3 dynamics under MR extends to normal physiology and under a longer-term alteration in nutrient availability that has beneficial health effects, we investigated the effects of MR on H3K4me3 *in vivo* under conditions previously shown to alter methionine metabolism, improve metabolic physiology in liver, extend life-span and reduce global levels of H3K4me3^32,45–47^. Thus, this mouse model of liver physiology in combination with the human cancer cell model provides a breadth of model systems covering both short- and long-term, both *in vitro* and *in vivo*, and both pathological and healthy conditions. We focused on liver in profiling the epigenomics and transcriptomics because it is the metabolic organ which is most responsive to metabolic reprogramming and there are liver-related phenotypes associated with methionine restriction related to metabolic health^46^. Livers were obtained from C57BL6 adult mice on a diet with either high methionine (0.84% w/w) or low, methionine-restricted (MR) methionine (0.12% w/w) for 12 weeks, and ChIP-seq of H3K4me3 was conducted (Fig 1F). Quantitation of bulk levels of H3K4me3 under these two conditions confirmed as was previously published that this level of dietary methionine restriction was sufficient to induce reduction of H3K4me3^32^. Physiological MR also revealed that the overall distribution of H3K4me3 is conserved between the different nutrient conditions as corroborated by conserved peak numbers (CV = 0.06, Fig S2A), location (Jaccard index = 0.82, Fig S2B) and marked genes (Jaccard index = 0.90, Fig S2C). High correlations in the values of peak height, area and width between high methionine and low methionine conditions were also observed with peak width as the most conserved (Fig 1G, linear regression coefficient = 0.99 for width compared to 0.85 for height and 0.80 for area). Consistent with what was found in human cancer cells, H3K4me3 peaks gained or lost in response to MR in mouse liver also tended to be significantly smaller peaks than those conserved in both high and low methionine conditions (Wilcoxon rank-sum P-value<1e-3 for 5 out of the 6 comparisons, Fig S2D, S2E) and were not enriched with specific biological function compared to randomly chosen genes containing H3K4me3 (median(-log_10_(enrichment Q-value)) = 3.09 for gained peaks compared to 4.03, 4.76 and 3.39 for three random gene sets with identical size, Fig S2F, and median(-log_10_(enrichment Q-value)) = 14.62 for lost peaks compared to 16.58, 15.75 and 13.27 for three random gene sets with identical size, Fig S2G). Reduction in the overall H3K4me3 signal was also observed (Fig S2H, S2I) as was a conserved distribution of genomic elements (84% promoter peaks and 16% non-promoter peaks, Fig S2J).

To assess the robustness of the quantitative peak descriptors to variation in ChIP-seq data analysis methodologies, we repeated the calculations using several different peak-calling algorithms. Although the total number of peaks (CV = 0.12 for human cancer cells, Fig S3A, and CV = 0.24 for mouse liver, Fig S3B) and total length of genomic regions covered by peaks (Jaccard index = 0.60 for human cancer cells, Fig S3C, and Jaccard index = 0.43 for mouse liver, Fig S3D) exhibited some variation among methods, reads mapped to peaks (CV = 0.0073 for human cancer cells, Fig S3E, and CV = 0.0078 for mouse liver, Fig S3F), genes associated with peaks (Jaccard index = 0.78 for human cancer cells, Fig S3G, and Jaccard index = 0.66 for mouse liver, Fig S3H), and dynamics of peak height, area and width (average Spearman’s rank correlation coefficient = 0.92 for human cancer cells and 0.86 for mouse liver, random permutation test P-values < 1e-323 for all comparisons, Fig S4) were concordant, implying that quantitation of peak dynamics is robust to the ChIP-seq data analysis method used. Taken together, these results indicate that changes in metabolism mediated by methionine availability, despite altering global levels of histone methylation, do not induce a complete redistribution of H3K4me3 marks on the genome.

### H3K4me3 peak width dynamics encode biological information

Although the genomic positioning of H3K4me3 was conserved in response to changes in methionine availability, we observed that peak height, area, and width were affected to different extents. To further investigate these differences, we computed the Spearman’s rank correlation coefficients between fold changes in peak height, area and width for both HCT116 cells and mouse liver and found that fold changes of width and the other two parameters were less correlated (Spearman’s rank correlation coefficient < 0.4) in both models (Fig 2A, 2B) relative to other comparisons. The lower correlation between changes in peak width and changes in the other two peak features suggested that a change in peak width might encode a different dimension of information. To test this hypothesis, we conducted gene set enrichment analysis (GSEA) on the peaks with the most sensitive (500 largest fold change) and robust (500 smallest fold change) height, area and width (Fig 2C) using the Molecular Signatures Database (MSigDB)^48^. For each subset of peaks, the distribution of the enrichment Q-values of the top 100 enriched (One-sided Fisher’s exact Q-value < 0.05) pathways was used to quantify association of this peak subset with biological functions. Three random peak subsets of equivalent size were also generated for comparison. In cultured cancer cells, the change in peak width exhibited the strongest and only significant signal for enrichment of specific biological processes as measured by the overall distribution of the resulting Q-values (Wilcoxon rank-sum P-value and Kolmogorov-Smirnov P-value < 1e-17 for comparisons between sensitive width and all other peak subsets, Fig 2D, S5A). In mouse liver, the consistency of the peak width exhibited the only signal above the randomly chosen sets of H3K4me3 that determined the background (Wilcoxon rank-sum P-value and Kolmogorov-Smirnov P-value < 1e-6 for comparisons between robust width and all other peak subsets, Fig 2E, S5B). This enrichment for biological function was found to be robust with respect to the size of the peak sets used (Fig S6A, S6B). This surprising finding appears to indicate that biological information that occurs in response to changes in methionine availability is encoded in H3K4me3 peak width.

**Figure 2.**
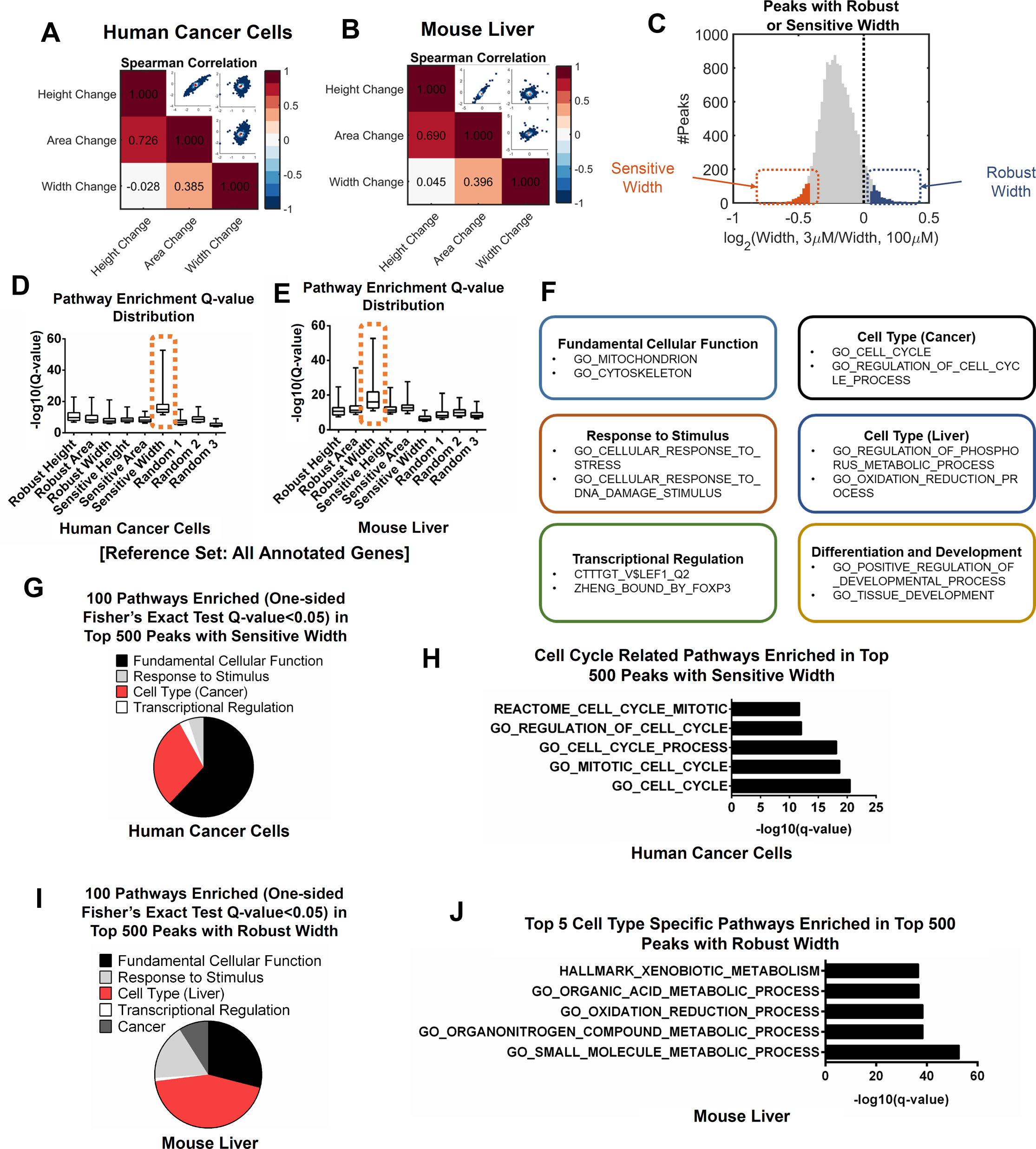
H3K4me3 peak width dynamics encode biological information. A. Heat map showing Spearman’s rank correlation coefficients among fold changes of H3K4me3 peak heights, areas and widths under MR in human cancer cells. The upper right part shows density scatter plots comparing the corresponding fold change values. B. Same as in (A) but for mouse liver. C. Distribution of fold changes (defined by the values in low methionine condition divided by the corresponding values in high methionine condition) in peak width in all peaks (grey), top 500 peaks with sensitive width (orange) and top 500 peaks with robust width (blue) in human cancer cells. D. Distributions of MSigDB pathway enrichment Q-values in peak sets with different dynamics under MR in human cancer cells. E. Same as in (D) but for mouse liver. F. Examples of annotated pathways in each category. G. Annotation of top 100 MSigDB pathways enriched in 500 H3K4me3 peaks with sensitive width in human cancer cells. H. Cell cycle related pathways enriched in 500 H3K4me3 peaks with sensitive width in human cancer cells. I. Same as in (G) but for robust width in mouse liver. J. Top 5 liver specific pathways enriched in 500 H3K4me3 peaks with robust width in mouse liver.

To further quantify the information contained in peak width, we annotated the top 100 enriched pathways in each case by classifying them into categorical subsets involving fundamental cellular function, response to stimulus, cell type (cancer, proliferation related pathways for human cancer cells and metabolism, multicellular organ related pathways for mouse liver), transcriptional regulation, differentiation and development, and cancer (Fig 2F). We next computed the fraction of each category in the top 100 pathways enriched for the 500 peaks with the most sensitive width in cancer cells and with the most robust width in liver. Unexpectedly, these two sets were enriched with cell type specific biological functions. Proliferation and cancer related pathways were enriched in peaks with sensitive width in cancer cells (30 out of the top 100 pathways, Fig 2G, 2H), while metabolism and multicellular organ related pathways were enriched in peaks with robust width in mouse liver (44 out of the top 100 pathways, Fig 2I, 2J). Importantly, the increased proportion of cell type specific functions in these peaks did not appear to a similar extent in randomly chosen genes with H3K4me3 marks (6 out of the top 100 pathways on average for human cancer cells, Fig S6C, and 8 out of the top 100 pathways on average for mouse liver, Fig S6D). Furthermore, these observed patterns of enrichment were preserved when using the set of all genes with H3K4me3 as the reference gene set instead of the complete set of annotated genes (Fig S7). Together these findings indicate that the peak width dynamics in response to MR encodes information about biological function.

### H3K4me3 peak width dynamics encode cell type specific transcription factor-binding preferences

To further explore this relationship, we probed additional aspects of genomic architecture. Using a motif analysis algorithm, HOMER^49^, we searched for transcription factor (TF) binding motifs enriched in the subsets of the different geometrical features of H3K4me3 peaks that change or remain consistent during MR. The complete set of H3K4me3 peaks was used as the background model (Fig 3A). In cultured cancer cells, TF binding motifs were only enriched (One-sided Fisher’s exact test Q-value < 0.05) in peaks with sensitive width (78 motifs enriched in sensitive width compared to 1 for robust area, 2 for robust width and 0 for all other peak sets, Fig 3B, S8A), while in mouse liver TF binding motifs were only enriched in peaks with robust width (249 motifs enriched in robust width compared to 0 for all other peak sets, Fig 3C, S8B). These findings were also found to be robust to the size of peak subsets chosen (Fig S8C, S8D). These two sets of TF binding motifs have little overlap (Jaccard index = 0.05 between the two sets of 10 motifs with smallest enrichment Q-values), suggesting that the role of H3K4me3 dynamics in relation to TF binding has a tissue specific function. The top scoring TF motifs in cancer cells that associated with sensitive H3K4me3 peak width dynamics tended to involve cancer-associated TFs such as c-MYC and ETS in cancer cells (Fig 3D) and liver-specific TFs such as RXR and ESRRB in liver (Fig 3E).

**Fig 3.**
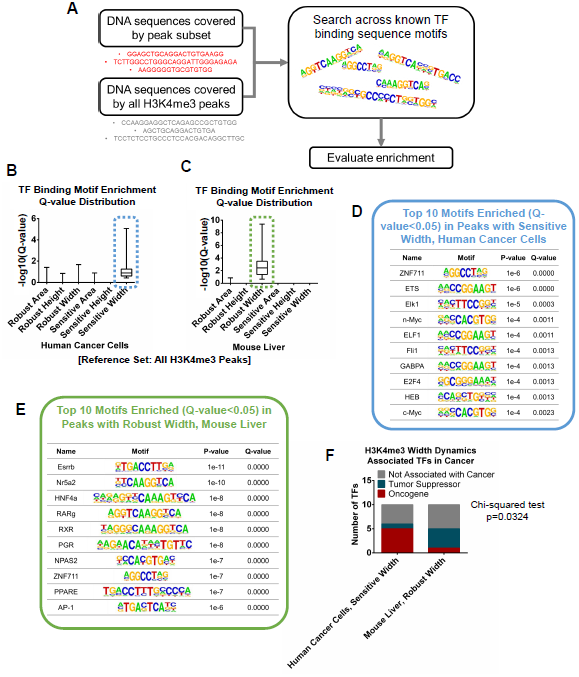
H3K4me3 peak width dynamics encode cell type specific transcription factor binding preferences. A. Framework for the TF binding motif enrichment analysis. B. Distributions of TF binding motif enrichment Q-values in H3K4me3 peak sets with different dynamics under MR in human cancer cells. C. Same as in (B) but for mouse liver. D. Top 10 TF binding motifs enriched in 500 H3K4me3 peaks with sensitive width in human cancer cells. E. Same as in (D) but for robust width in mouse liver. F. Number of oncogenes and tumor suppressors in top 10 TFs enriched in H3K4me3 peaks with sensitive width in human cancer cells and those with robust width in mouse liver.

To investigate the binding of these TFs, we obtained ChIP-seq datasets for two TFs putatively associated with H3K4me3 width in each system (Fig S9A) and computed the overlap between the TF binding sites and H3K4me3 peak subsets. We also chose one TF without binding motifs associated with peak width for each system as a negative control (Fig S9B). We found that the cancer-related TFs, MYC (Fig S9C) and ELF1 (Fig S9D), indeed had binding sites enriched (One-sided Fisher’s exact Q-value < 1e-10) in peaks with sensitive width in human cancer cells, while the liver-specific TFs, HNF4a (Fig S9E) and RXRa (Fig S9F-S9H), were also enriched (One-sided Fisher’s exact Q-value < 1e-10) in peaks with robust width in mouse liver. Such patterns were not observed in the TFs used as negative controls (One-sided Fisher’s exact Q-value = 1, Fig S9I, S9J). We also observed the strongest correlation between width dynamics and the number cell type specific TFs bound to the genomic region containing the H3K4me3 peak (One-way ANOVA P-value = 3.53e-319 for width changes in human cancer cells and 2.98e-189 for width changes in mouse liver, Fig S10). This concordance suggests that cell identity is encoded in the H3K4me3 width dynamics but not in any other structural or locational aspect of H3K4me3. A previous study has found that broad H3K4me3 peaks are associated with tumor suppressor gene function in normal cells^40^. We also found that TFs with binding motifs enriched in preserved H3K4me3 peak width in liver tended to be tumor suppressors, while those enriched in sensitive H3K4me3 peak width in cancer cells tended to be oncogenes (Chi-squared test P-value = 0.0324, Fig 3F), suggesting that H3K4me3 width dynamics under MR may also encode information about tumor suppression or progression, depending on the cell or tissue type. Altogether, these analyses highlight H3K4me3 peak width dynamics as the information carrier under MR in both human cancer cells and mouse liver.

### H3K4me3 peak width dynamics predict differential gene expression under MR

Despite a well-known association of the H3K4me3 mark with active promoters, the exact role of H3K4me3 in mediating gene expression is a matter of current debate^50,51^. We therefore asked whether any aspects of genomic alterations in H3K4me3 track with changes in gene expression under MR. We first quantified gene expression levels using RNA sequencing (RNA-seq) under high and low methionine conditions and correlated the gene expression levels with H3K4me3 peak features again in both cancer cells and mouse liver (Fig 4A). In cancer cells, we observed 7709 expressed genes and 5118 non-expressed genes marked by H3K4me3 (Fig S11A). There were also 11412 expressed genes without H3K4me3 marks, indicating in our models that the presence of the H3K4me3 mark is neither necessary nor sufficient for gene expression. Nevertheless, expressed genes with no H3K4me3 had significantly lower expression levels (Wilcoxon rank-sum P-value < 1e-323, Fig S11B), supporting the well-characterized association between H3K4me3 and active gene expression. This observation was further corroborated by significantly smaller H3K4me3 peak sizes in non-expressed genes (Wilcoxon rank-sum P-values < 1e-26 for height, area and width, Fig S11C-S11E). An evaluation of the correlation coefficients between H3K4me3 peak height, area, width and gene expression levels revealed significant positive correlations (Spearman’s rank correlation coefficient > 0.2, random permutation test P-value < 1e-50) between all three peak size descriptors and gene expression levels in both high and low methionine conditions (Fig 4B, S11F). Consistently, in mouse liver, we identified 9417 expressed and 2449 non-expressed genes with H3K4me3, as well as 9282 expressed genes without H3K4me3 (Fig S12A). In accordance with our findings in cultured human cells, strong correlations between the presence of H3K4me3 and gene expression levels were also observed in liver (P-value < 1e-20 for all Wilcoxon rank-sum and random permutation tests, Fig 4C, S12B-S12F). Thus, in our models, the presence of an H3K4me3 peak, while not a requirement for gene expression, predicts overall whether a gene is expressed, and the magnitude of the peak does appear to contain information about overall gene expression level. This finding is consistent with the well-known association between H3K4me3 around transcription start sites and active transcription^52^.

**Figure 4.**
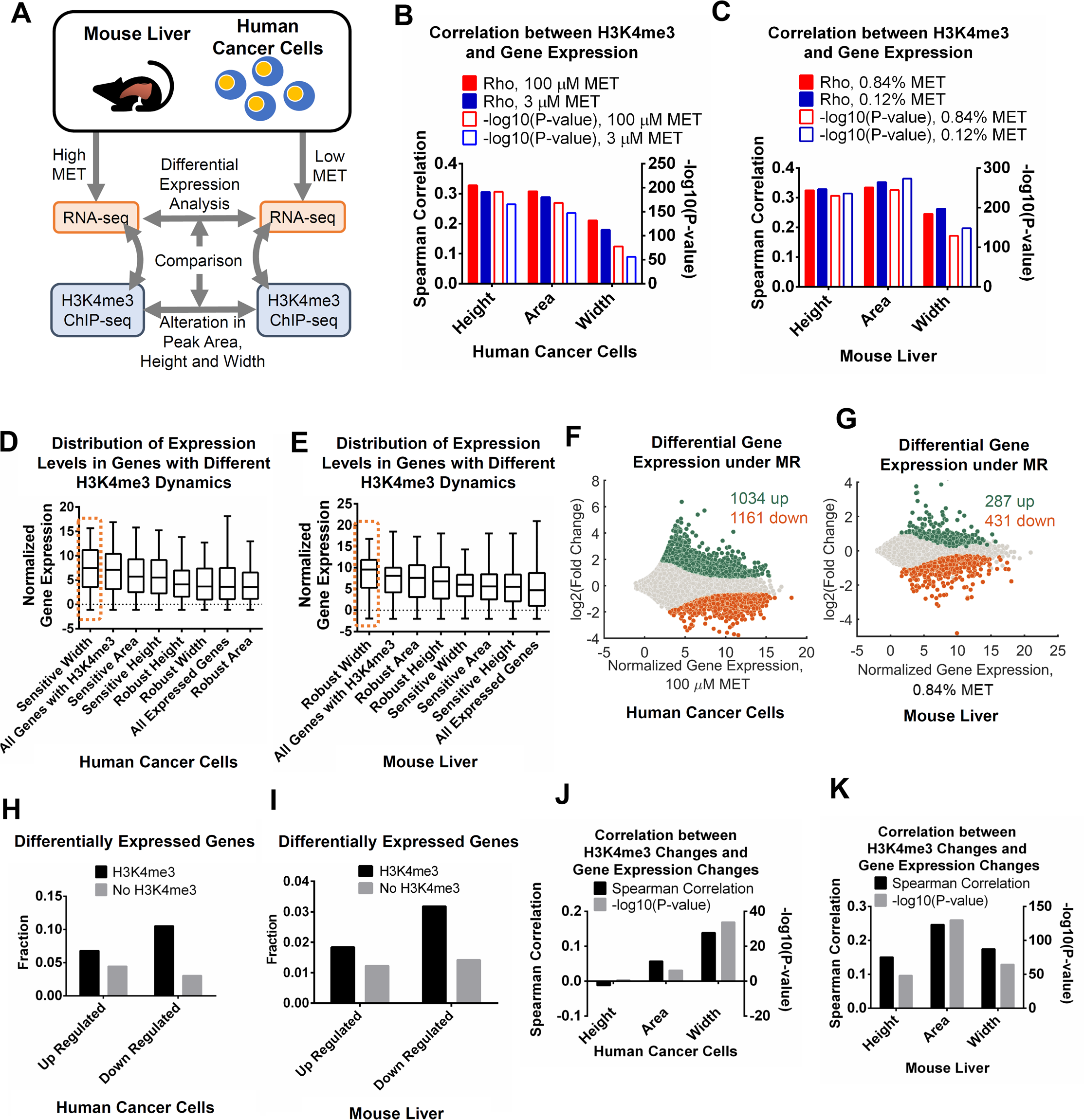
H3K4me3 peak width dynamics predict differential gene expression under MR. A. Framework of RNA-seq data analysis. B. Spearman correlation between peak size descriptors and gene expression levels in human cancer cells. C. Same as in (B) but for mouse liver. D. Expression levels of genes associated with different H3K4me3 dynamics under high methionine conditions in human cancer cells. The gene set with sensitive width which has been demonstrated to associate with cell type specific biological functions and TF binding is highlighted. E. Same as in (D) but for mouse liver. F. Differential gene expression in human cancer cells under MR. G. Same as in (F) but for mouse liver. H. Fraction of differentially expressed genes in genes with or without H3K4me3 in human cancer cells. I. Same as in (H) but for mouse liver. J. Spearman correlation between H3K4me3 changes and gene expression changes under MR in human cancer cells. K. Same as in (J) but for mouse liver.

Having established a baseline for the relationship between H3K4me3 and gene expression, we next sought to study whether changes in the geometrical features of H3K4me3 are connected to changes in gene expression. We compared expression levels of genes associated with different H3K4me3 dynamics in cancer cells (Fig 4D) and liver (Fig 4E) under high methionine conditions. In addition to our previous findings that sensitive H3K4me3 peak width in cancer cells and robust H3K4me3 peak width in mouse liver are indicative of biological function, we found that genes associated with these peaks also exhibit significantly higher expression levels (Wilcoxon rank-sum and Kolmogorov-Smirnov P-values < 0.05, Fig 4D, 4E, S13). We then conducted differential expression analysis to identify the genes with altered expression in response to MR. In human cancer cells, we found 1034 genes up-regulated (Wald test Q-value < 0.05, log_2_(fold change) > 0) and 1161 genes down-regulated (Wald test Q-value < 0.05, log_2_(fold change) < 0) under MR (Fig 4F, S14A, S14B) and 287 up-regulated (Wald test Q-value < 0.05, log_2_(fold change) > 0) and 431 down-regulated (Wald test Q-value < 0.05, log_2_(fold change) < 0) genes in mouse liver (Wald test Q-value < 0.05) (Fig 4G, S14C, S14D) in diverse classes of genes. It is noteworthy that differential gene expression *in vivo* is confounded by factors including composition of liver by different cell types and larger variation between individual mice. Thus, we expect less genes to be differentially expressed in mouse liver. We next asked if changes to or consistencies in the geometrical features of H3K4me3, especially in peak width, can predict changes in gene expression. We first compared the fraction of differentially expressed genes that contain or are absent of H3K4me3. We found that in both cancer cells and liver, H3K4me3-marked genes were enriched in both up- and down-regulated genes (One-sided Fisher’s exact P-value = 1.82e-12 for up-regulated genes and 1.89e-98 for down-regulated genes in human cancer cells, Fig 4H, and 4.22e-4 for up-regulated genes and 4.43e-16 for down-regulated genes in mouse liver, Fig 4I), suggesting that the presence of H3K4me3 notes a tendency of differential expression during MR. Next, we correlated fold changes in H3K4me3 peak height, area, width with changes in gene expression levels and found that H3K4me3 peak width dynamics significantly correlated (random permutation test P-value < 0.05, Spearman’s rank correlation coefficient > 0.1) with alterations in gene expression levels, and this correlation was consistent in both models (Spearman’s rank correlation coefficient = 0.14 for human cancer cells and 0.17 for mouse liver, Fig 4J, 4K). On the other hand, although H3K4me3 peak height and area dynamics were also found to correlate with differential gene expression in mouse liver (random permutation test P-value < 0.05, Spearman’s rank correlation coefficient > 0.15, Fig 4K), the strength of these correlations was smaller in human cancer cells (Spearman’s rank correlation coefficient < 0.06, Fig 4J). Restricting the analysis to the subset of peaks located at promoters (Fig S14E) or altering the data analysis pipeline (Fig S14F) had minimal effects on the resulting correlation coefficients (absolute differences in Spearman’s rank correlation coefficients < 0.04 for data in Fig 4J, Fig S14E, S14F). Interestingly, we also observed a stronger correlation between H3K4me3 width changes and gene expression changes in genes with H3K4me3 peaks bound by more TFs putatively associated with H3K4me3 width in both human cancer cells (Spearman’s rank correlation coefficient between width changes and gene expression changes = 0.19 in peaks bound by 2 TFs compared to 0.10 for 0 TF and 0.12 for 1 TF, Fig S14G) and mouse liver (Spearman’s rank correlation coefficient between width changes and gene expression changes = 0.22 in peaks bound by 2 TFs compared to 0.18 for 0 TF and 0.19 for 1 TF, Fig S14H), suggesting that TFs associated with H3K4me3 width dynamics regulate expression of their target genes thus mediating the connection between H3K4me3 dynamics and gene expression dynamics. Taken together, H3K4me3 width dynamics, but not area or height, is the predictor of alterations in gene expression in both human cancer cells and mouse liver. Moreover, peak width also encodes information of gene expression levels under normal methionine conditions in addition to cell type specific biological functions and TF binding preferences.

## Discussion

Numerous studies have shown that global levels of histone and DNA modifications are influenced by metabolism and changes to nutrient availability^1–5^. Biological outcomes however result from reprogramming of chromatin state which influences regulation of gene expression and these mechanisms are still poorly understood. To our knowledge, this study is the first to identify general principles about how specific aspects of the genomic architecture of a histone mark are affected by nutrient availability.

### H3K4me3 peak width encodes information about biological function and gene expression dynamics

We found that H3K4me3, a chromatin mark known to associate with active transcription, responds to MR with a global compression of peak area and height across most modified sites, which is consistent with the substantial reduction of bulk levels observed previously^6,32^. H3K4me3 peak width, despite being the most conserved feature under MR, uniquely encoded, in its dynamics, information about cell identity, TF binding preferences, tumor suppression and gene expression that are not reflected in changes in other aspects of H3K4me3. Peak width dynamics was also the only predictor of gene expression alterations in both human cancer cells and mouse liver. In addition to peak width encoding information about cell identity^39,40^, we showed that its dynamics under alterations in methionine metabolism is also a link between H3K4me3 and changes to gene expression. This finding extends our understanding of how metabolism influences chromatin biology since it identifies novel aspects of biological specificity in H3K4me3 dynamics across the genome.

Finally, we observed opposing effects in cultured human cancer cells and in mouse liver. H3K4me3 width changes in cancer cells and width conservation in liver marked biological functions and gene expression changes. In both cases, the biological associations with these dynamics may be attributed to the function of the tissue and the physiological or pathophysiological status of the model in relation to the ongoing gene expression programs. The discrepancy between cancer cells and healthy tissue may also reflect differences between cancerous and normal cell types in responding to alterations in environmental factors. We speculate that, at least in the context of MR, cell type specific functions in normal tissues are more robust under alteration in environmental variables to ensure normal function, while cancer cells have more flexibility which potentially maximizes fitness in a varying environment. Notably, we found that genes related to proliferation and survival of HCT116 cancer cells identified in a CRISPR screen^53^ exhibited significantly elevated sensitivity in both H3K4me3 width (Wilcoxon rank-sum P-value = 1.47e-136) and gene expression (Wilcoxon rank-sum P-value = 8.15e-116) in response to MR (Fig S15), suggesting that the influences of MR on H3K4me3 peak width are indeed linked to functional outcome. Although further investigation is needed to unravel the mechanism conferring this profound discrepancy between pathological and healthy models, the general principle that the peak width is the most informative parameter in H3K4me3 dynamics upon changes in nutrient availability is conserved in both models.

### Passive and active roles for H3K4me3

Although long appreciated to be a signature of active gene expression^52^, the role of H3K4me3 in regulation of gene expression remains controversial^51^. There is evidence that H3K4me3 interacts with transcriptional and splicing machineries to regulate gene expression^54,55^, while other studies have concluded that the timing of H3K4me3 changes across biological conditions precludes it having an active role in gene expression^56,57^. Other studies have concluded that the mark may affect the robustness of transcriptional programs^39^. Recent studies of H3K4me3 during early embryo development have further suggested that the function of H3K4me3 in this process may be to counteract DNA methylation at specific genomic regions^41,43^, supporting a function independent of facilitating transcription. In this study, we showed that reprogramming of the H3K4me3 landscape under an alteration in nutrient availability was indeed related to alterations in gene expression but through a specific mechanism involving the dynamics of H3K4me3 peak width. Although more work is needed to establish definitive causality between H3K4me3 and gene expression dynamics, our studies support a model that a subset of overall gene expression appears to be responsive to changes in H3K4me3 and that these programs are encoded in changes in peak width.

## Summary

There remain numerous unanswered questions on the interaction between metabolism and chromatin biology. Chromatin status is a manifestation of more than 100 covalent modifications and multiple assembly factors^58–60^ and numerous metabolic pathways beyond one-carbon metabolism directly interact with chromatin. It is to be determined how general are the principles we found regarding H3K4me3 peak width and whether they extend to other marks or to other metabolic changes that alter methylation such as those regulated by mitochondrial metabolism and alpha-ketoglutarate. For example, H3K4me3 levels are also reduced by knockdown of the histone methyltransferase MLL1 in HCT116 cells^61^, but the conserved and unique features in this process compared to MR is not clear, although there is existing literature that has defined some aspects of the specificity of the requirements of methyltransferases^62–65^. In conclusion, this study unravels simple yet fundamental principles of how metabolism influences chromatin biology, that is, almost all aspects of H3K4me3 biology, including cell identity, tumor suppression and progression and gene expression are encoded in H3K4me3 width dynamics.

## Materials and Methods

### Methionine restriction in human cancer cells and mouse

HCT116 cells were cultured as previously described ^32^. Briefly, HCT116 cells were obtained from ATCC and the stock used for this study was recently validated as bona fide HCT116 cells via the Duke University DNA Analysis Facility Human cell line authentication service. Cells were cultured in RPMI with 10% FBS (containing 30 micromolar methionine). Plated cells were first cultured in 100 micromolar methionine and switched to 3 micromolar for 24 hours upon harvesting lysates. Mouse feeding and tissue collection were performed as previously described^32^. Briefly, C57BL/6J mice were fed with either 0.84% (w/w) methionine control diet or 0.12% (w/w) methionine MR diet for 12 weeks and fasted before being sacrificed. Input chromatin was pooled from all samples in each replicate.

### H3K4me3 ChIP-seq data analysis

ChIP was performed as previously described^32^. In addition, we applied a spike-in normalization strategy for generating quantitative ChIP-seq data^66^, in which spike-in Drosophila chromatin and spike-in antibody for Drosophila H2Av (Active Motif cat#61686) was mixed with chromatin from the HCT116 cells before the chromatin IP step with a fixed ratio (2:1, Drosophila:HCT116 chromatin) as a reference. Libraries were prepared according to Illumina instructions and sequenced on the Illumina HiSEQ 2500 sequencer in Rapid Run Mode at the Duke GCB Sequencing Shared Resource. Reads were aligned to a combinational genome consisting of the human reference genome hg19 and the Drosophila reference genome dm6 using Bowtie2^67^. Alignment files were then filtered according to their alignment scores and down-sampled to ensure each file contains the same amount of unique Drosophila reads. Finally, reads mapped to hg19 in the normalized alignment files were kept for peak-calling and other following analyses. Replicate 1 for high methionine condition was not used due to abnormally high fraction of Drosophila-originated reads in this sample. Processing of alignment files was done using SAMtools^68^. Peaks were called using the --broad mode of MACS2^69^ and filtered with the criterion that enrichment Q-value is smaller than 10^−5^. For mouse liver, ChIP-seq reads were aligned to the mouse reference genome mm8 and normalized by sequencing depth.

### Peaks annotation and size descriptors computation

H3K4me3 peak regions called for different methionine conditions and replicates were merged using the bedtools merge command in BEDTools package^70^ to generate a combinational peak set for following annotation and computation of peak descriptors. Peaks were assigned to genes with transcription start site (TSS) closest to center of the peak region using Homer^49^. Peak height, area and width were computed on this combinational peak set using the extended reads coverage files (that is, number of fragments extended from ChIP-seq reads mapped to each base pair on the genome) generated by MACS2. Height was computed by searching for position with highest read coverage. Area was computed by integrating the reads coverage file over the peak region. Width was quantified by first normalizing reads coverage profile of each peak to a probability distribution function (*i.e*. area under the coverage curve = 1) and then calculating standard deviation accordingly. Percentage peak areas on each type of genomic elements were computed using C++ codes with genomic element annotation files in the HOMER package and reads coverage files generated by MACS2 as inputs. Replicates were merged by computing average between them.

### Comparison between peak calling methods

H3K4me3 peaks were called using Bayesian Change Point (BCP)^71^, MUltiScale enrIchment Calling for ChIP-seq (MUSIC)^72^ and MACS2 either with ( MACS2.broad) or without (iMACS2.narrow) the --broad option. Default parameters for each of these methods were used. Generation of unions and intersections of the peak sets and quantification of reads mapped to peak regions was conducted with BEDTools using the commands bedtools merge, bedtools intersect and bedtools coverage. Extended read coverage profiles were generated using MACS2 as described previously. Raw read coverage profiles were generated from the normalized alignment files using the command bedtools genomecov in BEDTools without applying the model in MACS2 for extension of reads to whole fragments. For the comparison of H3K4me3 changes between peak-calling algorithms, peaks called by the two algorithms assigned to the same gene were compared with each other.

### Pathway and TF binding motif enrichment analysis

Pathway enrichment analysis with all annotated genes as reference set was conducted using the Molecular Signatures Database (MSigDB) 6.0 (http://software.broadinstitute.org/gsea/msigdb/index.jsp) with gene lists associated with peak subsets as inputs. Gene sets H (hallmark gene sets), C1 (positional gene sets), CP (canonical pathways), CP:BIOCARTA (BioCarta gene sets), CP:KEGG (KEGG gene sets), CP:REACTOME (Reactome gene sets), C3 (motif gene sets), C4 (computational gene sets) and C5 (GO gene sets) were included in the analysis. For GSEA using all genes with H3K4me3 as the reference set, a complete list of pathways was first downloaded from MSigDB, then modified by removing all genes without H3K4me3. Enrichment P-values were then computed using a one-sided Fisher’s exact test and corrected using the Benjamini-Hochberg procedure. TF binding motif enrichment analysis was done by the module findMotifsGenome.pl in HOMER. Lists of tumor suppressors and oncogenes were obtained from the databases TSGene 2.0^73^ and ONGene^74^.

### TF ChIP-seq data analysis

Raw read files in SRA or FASTQ format for the TF ChIP-seq experiments were downloaded using the accession numbers in Fig S9A and Fig S9B, aligned to the reference genome hg19 for HCT116 cells and mm8 for mouse liver using Bowtie2, and filtered according to alignment scores using SAMtools. SRA files were converted to FASTQ format using the command fastq-dump in the SRA Toolkit (https://github.com/ncbi/sra-tools) before the alignment. Peaks were called using MACS2 without the --broad option and filtered with the criterion that the enrichment Q-value is smaller than 10^−5^. H3K4me3 peaks bound by a TF were defined as those H3K4me3 peaks overlapping with peaks called from the corresponding TF ChIP-seq data. Fold enrichment of TF binding in a H3K4me3 peak subset was defined as the ratio of fraction of peaks bound by the TF in this H3K4me3 peak subset relative to the fraction of peaks bound by the TF in the complete set of H3K4me3 peaks, that is:

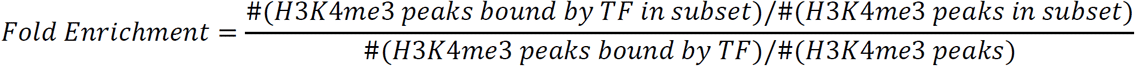

Enrichment Q-values were computed by correcting P-values from one-sided Fisher’s exact test using Benjamini-Hochberg procedure.

### RNA-seq

Total RNA from HCT116 cells and mouse liver under high and low methionine conditions was extracted using the PARIS kit (Life Technologies, cat#AM1921), polyA selected and then sent to the Weill-Cornell Epigenomics core (HCT116) and the Duke GCB Sequencing Shared Resource (mouse liver) for library preparation and sequencing. Libraries were sequenced either on the Illumina HiSEQ 2500 sequencer in Rapid Run Mode (HCT116) or on the Illumina HiSEQ 4000 sequencer (mouse liver).

### RNA-seq data analysis

Raw reads were aligned to the human reference genome hg19 and mouse reference genome mm8, respectively, using TopHat2^75^. Number of reads mapped to each gene feature was first quantified using HTSeq^76^ with the input of GTF files obtained from the UCSC Table Browser and then normalized using DESeq2^77^. Differential expression analysis was done with DESeq2. P-values were adjusted using the Benjamini-Hochberg procedure. Genes with adjusted differential expression P-values smaller than 0.05 were considered as differentially expressed.

### Fitness genes in human cancer cells

The list of fitness genes was extracted from Table S2 of the published study on CRISPR-based screen of proliferation and survival related genes in HCT116 cells^53^.

### Data visualization

Heatmaps and average profiles showing the H3K4me3 ChIP-seq signal were generated with the commands plotHeatmap and plotProfile in deepTools. Other heatmaps and density scatter plots were created using MATLAB. All bar graphs, pie graphs and box plots were created using GraphPad Prism. ChIP-seq tracks were created using Integrative Genomics Viewer (IGV)^78^.

### Codes and data availability

Codes and processed data are available at GitHub page of the Locasale Lab (https://github.com/LocasaleLab/H3K4me3_MET_Restriction). Raw data are available at the Gene Expression Omnibus (GEO) database with accession number GEO: GSE103602. Peak height, area, width and gene expression values in different conditions are available in Table S1.

## Acknowledgements

Support from the American Cancer Society (RSG-16-214-01-TBE) and National Institutes of Health (R01CA193256, R00CA168997, P30CA014236) are gratefully acknowledged. Z.D. thanks Dr. Ning Yin for helpful discussions. We thank Dwight Mattocks (Orentreich Foundation for the Advancement of Science) for assistance with animal husbandry.

## Author Contributions

Z.D. and J.W.L. designed the study and wrote the manuscript with input from S.J.M.; Z.D. performed the data analysis; S.J.M. performed the experiments with help from X.G. S.N.N. performed mice feeding and tissue harvesting.

**Figure S1.**
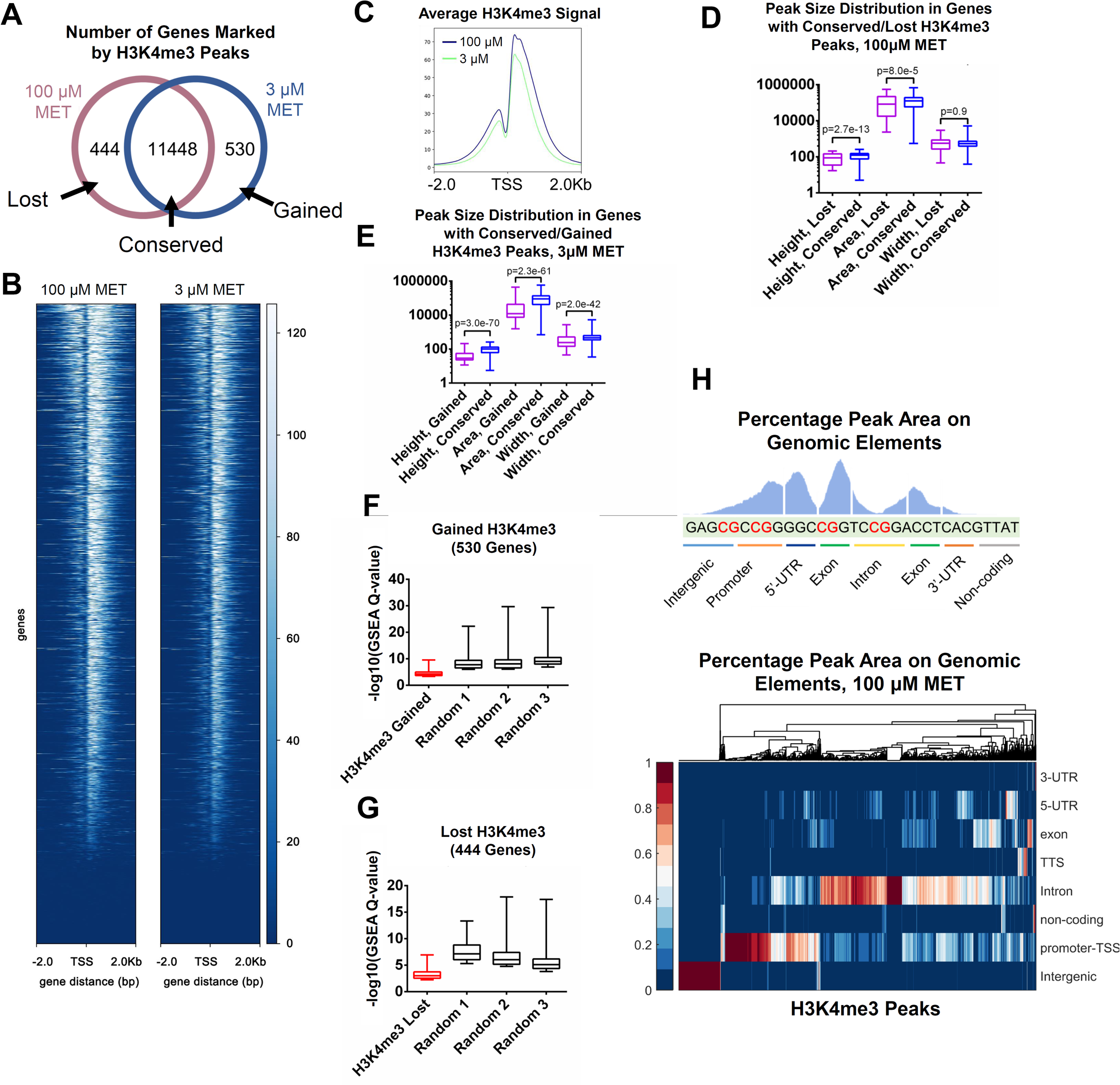
(Related to Fig 1) A. Number of genes marked by H3K4me3 peaks under high and low methionine conditions in human cancer cells. B. Average H3K4me3 profiles around transcription start sites (TSS)s under high and low methionine conditions in human cancer cells. C. Heatmap showing H3K4me3 signal around TSS of each marked gene under high and low methionine conditions in human cancer cells. D. Distribution of peak height, area and width under high methionine condition in genes with conserved or lost H3K4me3 peaks in human cancer cells. The P-values were computed from the Wilcoxon rank-sum test. E. Distribution of peak height, area and width under low methionine condition in genes with conserved or gained H3K4me3 peaks in human cancer cells. The P-values were computed from the Wilcoxon rank-sum test. F. Distributions of pathway enrichment Q-values in 530 genes with gained H3K4me3 and random gene sets with identical size in human cancer cells. G. Same as in (F) but for 444 genes with loss of H3K4me3. H. Definition of percentage peak area on genomic elements and its distribution under high methionine condition in human cancer cells.

**Figure S2.**
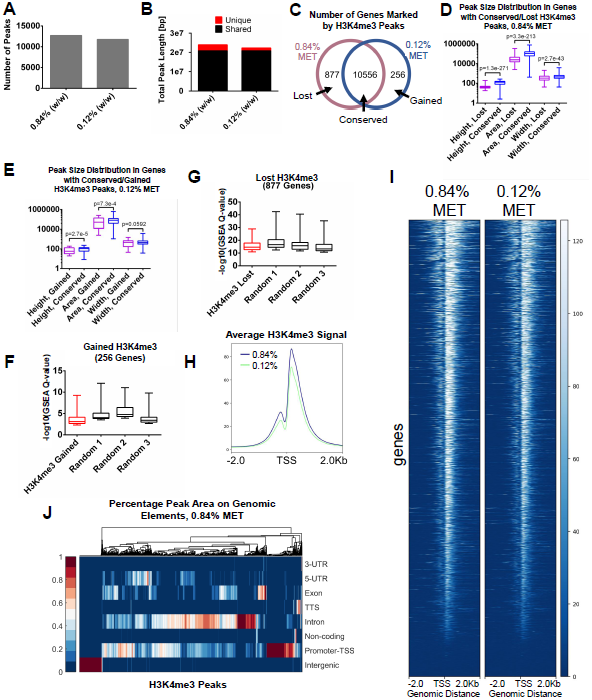
(Related to Fig 1) A. Shared and unique length of peak regions under high and low methionine conditions in mouse liver. B. Number of peaks called under high and low methionine conditions in mouse liver. C. Number of genes marked by H3K4me3 peaks under high and low methionine conditions in mouse liver. D. Distribution of peak height, area and width under high methionine condition in genes with conserved or lost H3K4me3 peaks in mouse liver. The P-values were computed from the Wilcoxon rank-sum test. E. Distribution of peak height, area and width under low methionine condition in genes with conserved or gained H3K4me3 peaks in mouse liver. The P-values were computed from the Wilcoxon rank-sum test. F. Distribution of pathway enrichment Q-values in 256 genes with gained H3K4me3 and random gene sets with identical size in mouse liver. G. Same as in (F) but for 877 genes with loss of H3K4me3. H. Average H3K4me3 profiles around TSSs under high and low methionine conditions in mouse liver. I. Heatmap showing H3K4me3 signal around TSS of each marked gene under high and low methionine conditions in mouse liver. J. Percentage peak area on genomic elements for each H3K4me3 peak under high methionine condition in mouse liver.

**Figure S3.**
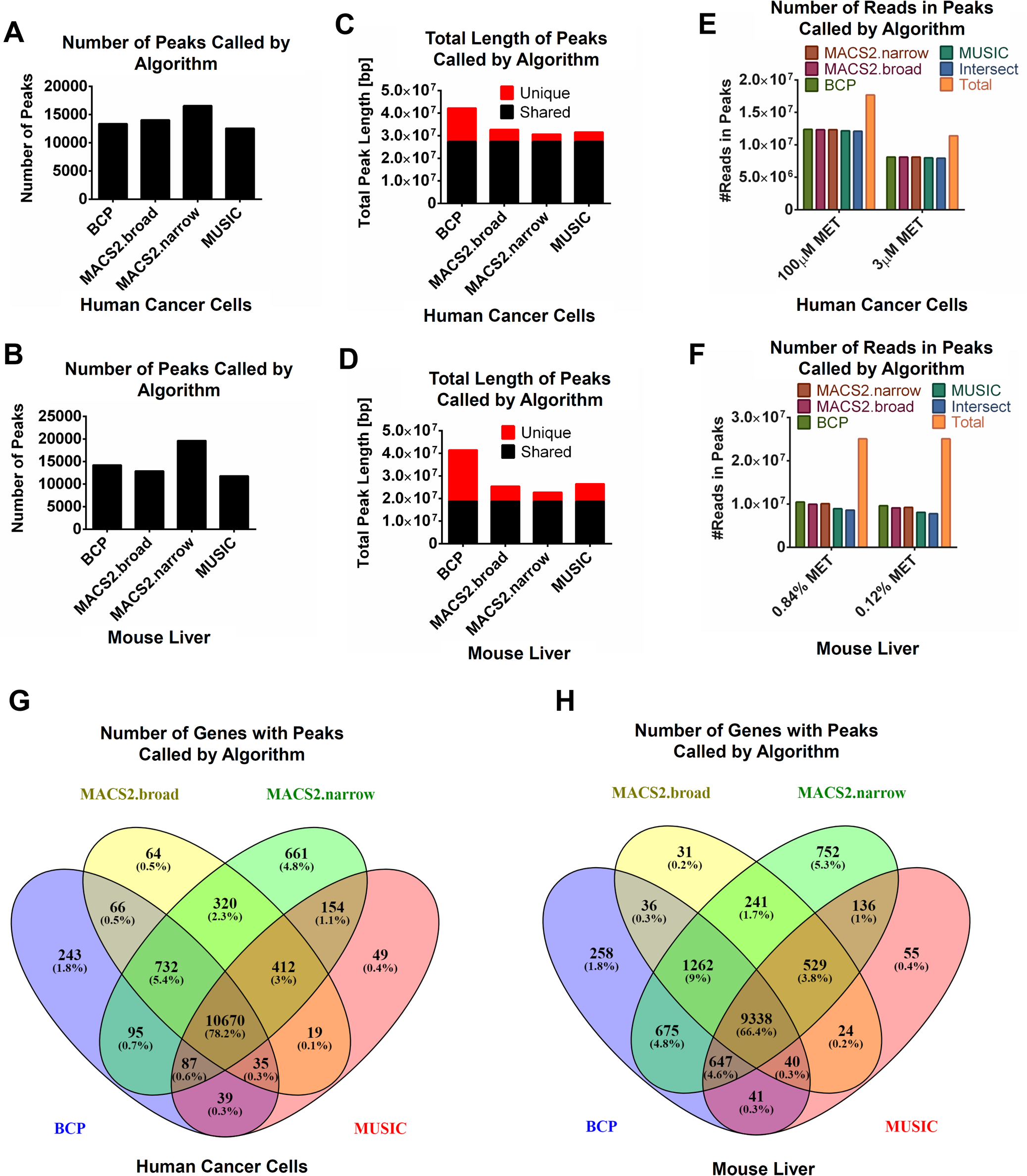
(Related to Fig 1) A. Number of peaks called by different peak-calling algorithms in human cancer cells. B. Same as in (A) but for mouse liver. C. Total length of peaks called by different peak-calling algorithms in human cancer cells. D. Same as in (C) but for mouse liver. E. Number of reads in peaks called by different peak-calling algorithms in human cancer cells. F. Same as in (E) but for mouse liver. G. Venn diagram of genes marked by peaks called by different peak-calling algorithms in human cancer cells. H. Same as in (G) but for mouse liver.

**Figure S4.**
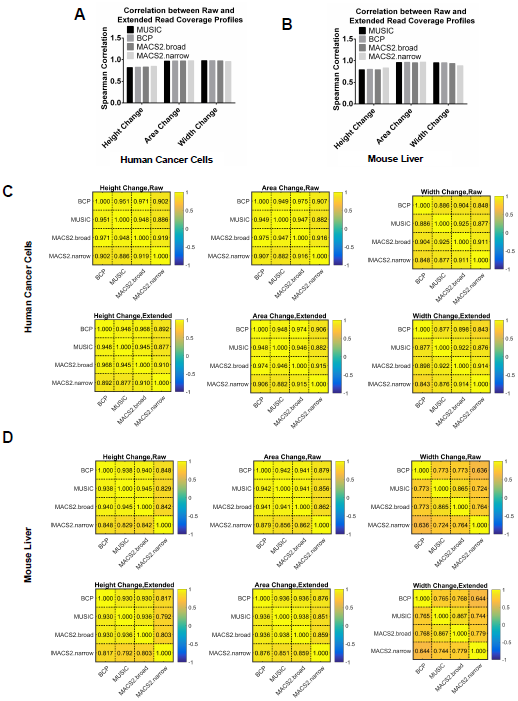
(Related to Fig 1) A. Spearman correlation between raw (i.e. without extension of reads to the whole fragment) and extended (i.e. with extension of reads to the whole fragment) read coverage profiles for changes in height, area and width in human cancer cells. B. Same as in (A) but for mouse liver. C. Spearman correlation between peak calling algorithms for changes in height, area and width in human cancer cells. D. Same as in (C) but for mouse liver.

**Figure S5.**
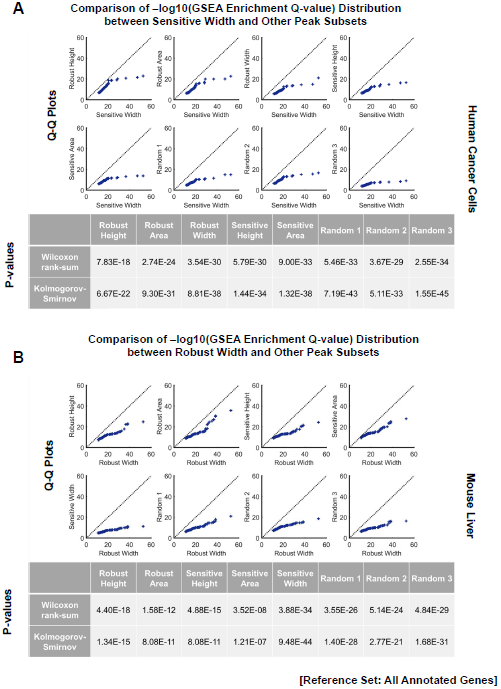
(Related to Fig 2) A. Quantile-Quantile (Q-Q) plots, Wilcoxon rank-sum P-values and Kolmogorov-Smirnov P-values comparing distributions of pathway enrichment Q-values in other peak subsets to the peak subset with sensitive width in human cancer cells. B. Same as in (A) but for robust width in mouse liver.

**Figure S6.**
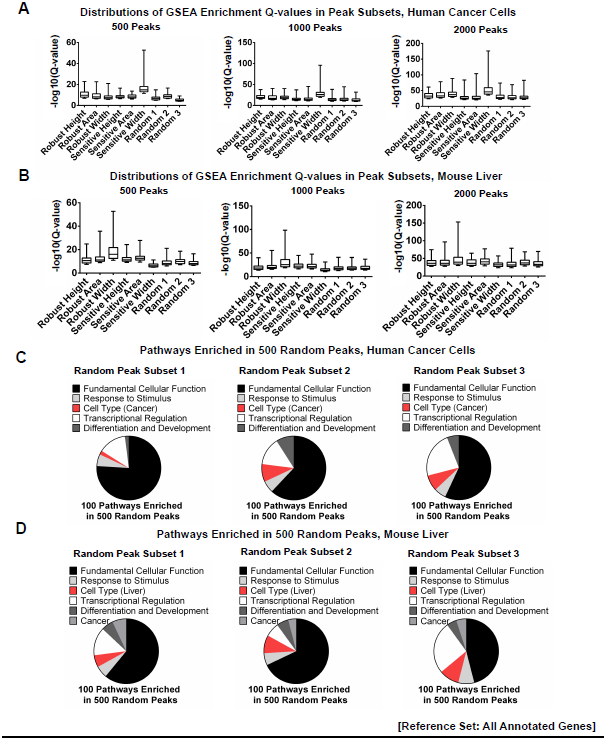
(Related to Fig 2) A. Pathway enrichment Q-value distributions in 500, 1000 and 2000 H3K4me3 peaks with different dynamics under MR in human cancer cells. B. Same as in (A) but for mouse liver. C. Annotation of top 100 enriched pathways in 3 sets of 500 random peaks in human cancer cells. D. Same as in (C) but for mouse liver.

**Figure S7.**
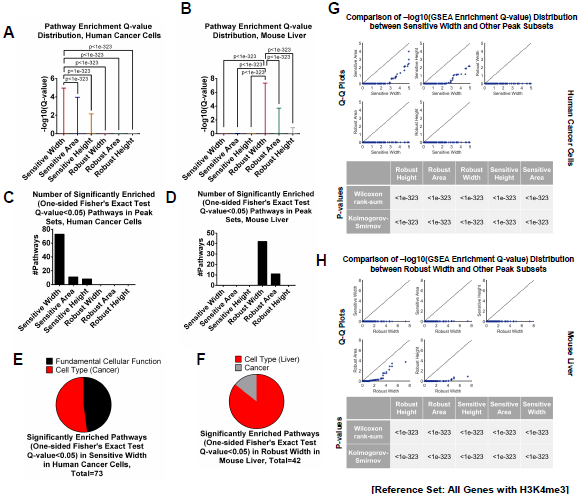
(Related to Fig 2) A. Distributions of pathway enrichment Q-values estimated using all genes with H3K4me3 as the reference set in human cancer cells. P-values were computed from Wilcoxon rank-sum test. B. Same as in (A) but for mouse liver. C. Number of significantly enriched pathways in peak subsets in human cancer cells. D. Same as in (C) but for mouse liver. E. Annotation of pathways significantly enriched in peak subset with sensitive width in human cancer cells. F. Same as in (E) but for robust width in mouse liver. G. Q-Q plots, Wilcoxon rank-sum P-values and Kolmogorov-Smirnov P-values comparing distributions of pathway enrichment Q-values in other peak subsets to the peak subset with sensitive width in human cancer cells. H. Same as in (G) but for robust width in mouse liver.

**Figure S8.**
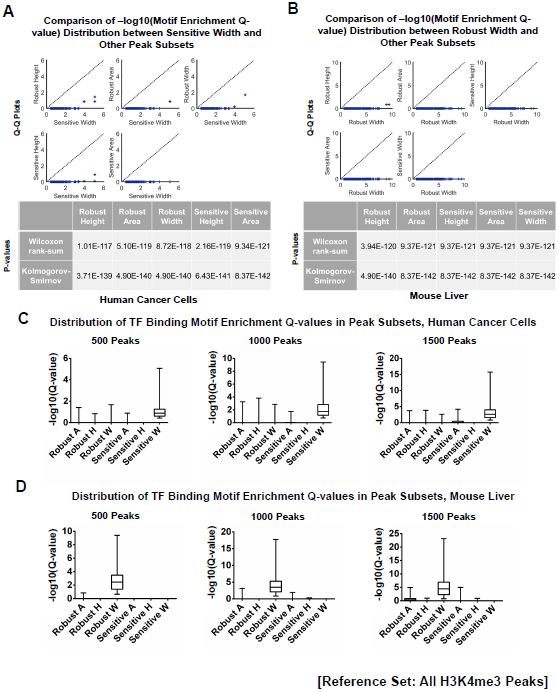
(Related to Fig 3) A. Q-Q plots, Wilcoxon rank-sum P-values and Kolmogorov-Smirnov P-values comparing distributions of TF binding motif enrichment Q-values in other peak subsets to the peak subset with sensitive width in human cancer cells. B. Same as in (A) but for robust width in mouse liver. C. TF binding motif enrichment Q-value distributions in 500, 1000 and 1500 H3K4me3 peaks with different dynamics under MR in human cancer cells. D. Same as in (C) but for mouse liver.

**Figure S9.**
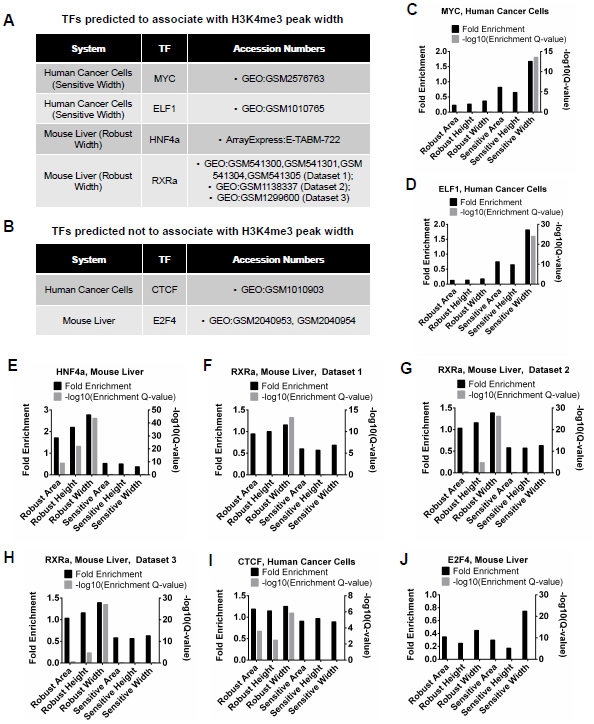
(Related to Fig 3) A. Table of ChIP-seq datasets used in the analysis of Transcription Factors (TF)s putatively associated with H3K4me3 peak width. Column 1 represents the system studies, Column 2 denotes the TF used, and Column 3 defines the accession numbers for the ChIP-seq data. B. Table of ChIP-seq datasets as in (A) but for TFs putatively not associated with H3K4me3 peak width. C. Fraction of peaks bound by MYC in a subset of peaks (defined on the x-axis) relative to the fraction of peaks bound by the TF in all H3K4me3 peaks (i.e. fold enrichment) in human cancer cells. Q-values represent an adjusted (Benjamin-Hochberg method) P-value obtained from a one-side Fisher’s exact test. Further information is contained in the Methods. D. Same as in (C) but for ELF1 in human cancer cells. E. Same as in (C) but for HNF4a in mouse liver. F. Same as in (C) but for RXRa in mouse liver based on Dataset 1. G. Same as in (C) but for RXRa in mouse liver based on Dataset 2. H. Same as in (C) but for RXRa in mouse liver based on Dataset 3. I. Same as in (C) but for CTCF in human cancer cells. J. Same as in (C) but for E2F4 in mouse liver.

**Figure S10.**
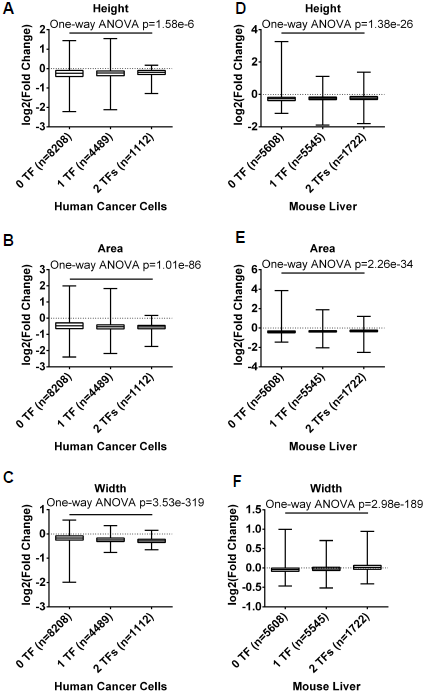
(Related to Fig 3) A. Distributions of H3K4me3 height changes (high and low methionine) in peaks bound by 0,1 or 2 TFs putatively associated with H3K4me3 width in human cancer cells. B. Same as in (A) but for area. C. Same as in (A) but for width. D. Distribution of H3K4me3 height changes in peaks bound by 0, 1, or 2 TFs putatively associated with H3K4me3 width in mouse liver. E. Same as in (D) but for area. F. Same as in (D) but for width.

**Figure S11.**
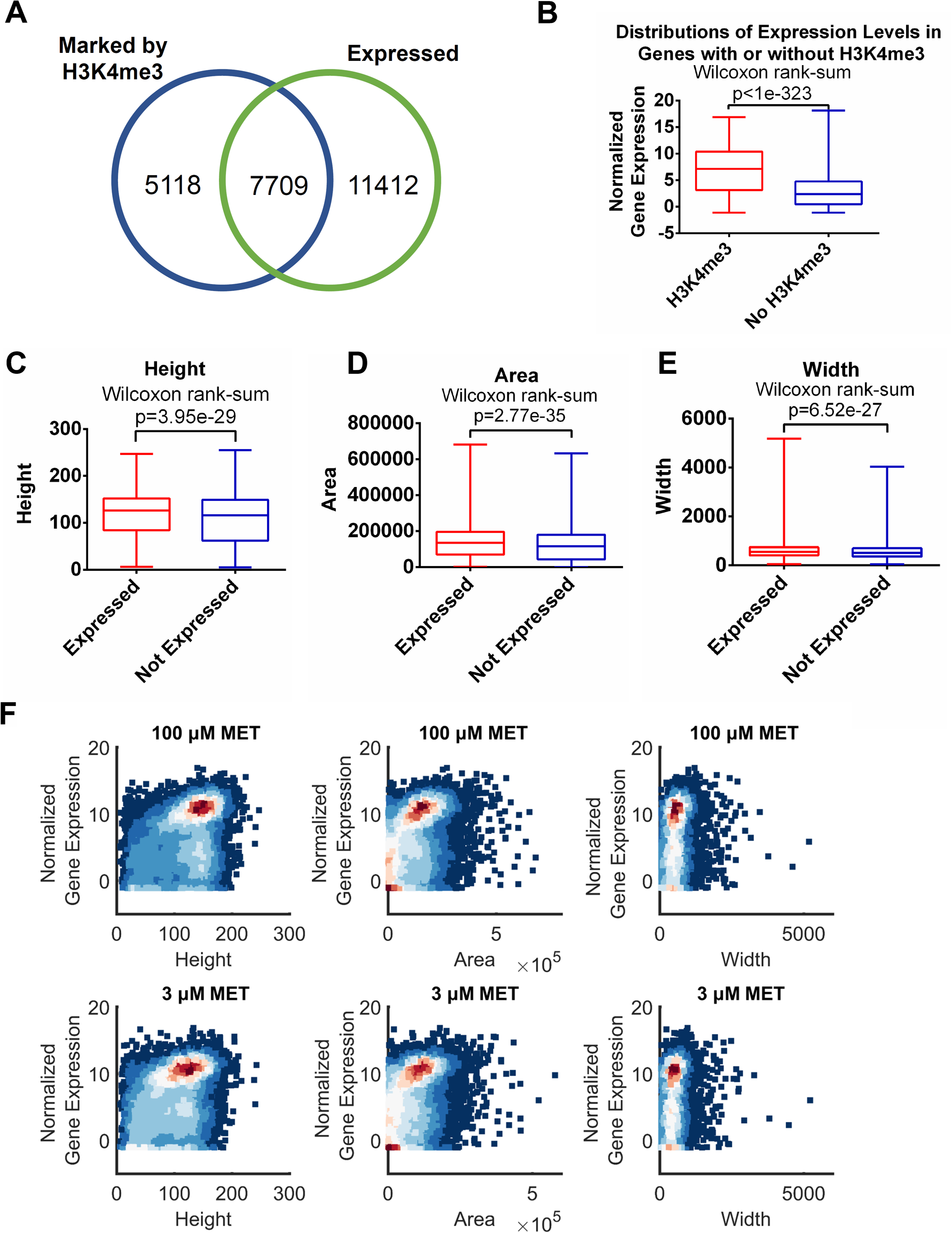
(Related to Fig 4) A. Number of genes marked by H3K4me3 peaks, expressed genes and their overlap in human cancer cells. B. Distribution of expression levels in genes with or without H3K4me3 in human cancer cells under high methionine conditions. C. Distribution of height in H3K4me3 peaks associated with expressed or non-expressed genes in human cancer cells under high methionine conditions. D. Same as in (C) but for area. E. Same as in (C) but for width. F. Density scatter plots comparing peak heights, areas and widths with gene expression levels under high and low methionine conditions in human cancer cells.

**Figure S12.**
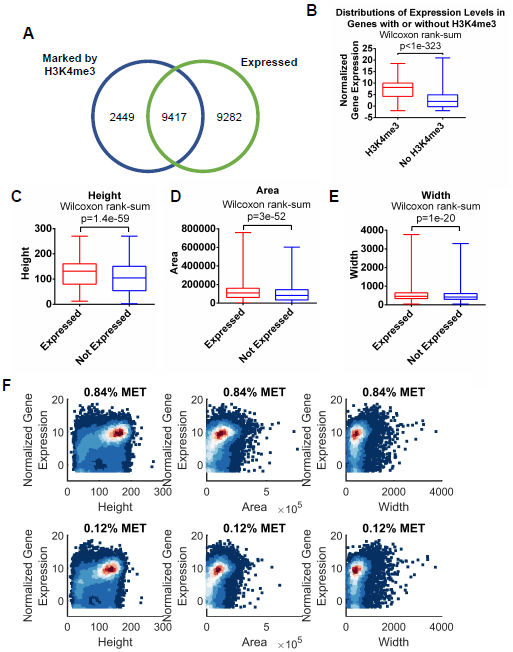
(Related to Fig 4) A. Number of genes marked by H3K4me3 peaks, expressed genes and their overlap in mouse liver. B. Distribution of expression levels in genes with or without H3K4me3 in mouse liver under high methionine conditions. C. Distribution of heights in H3K4me3 peaks associated with expressed or non-expressed genes in mouse liver under high methionine conditions. D. Same as in (C) but for area. E. Same as in (C) but for width. F. Density scatter plots comparing peak heights, areas and widths with gene expression levels under high and low methionine conditions in mouse liver.

**Figure S13.**
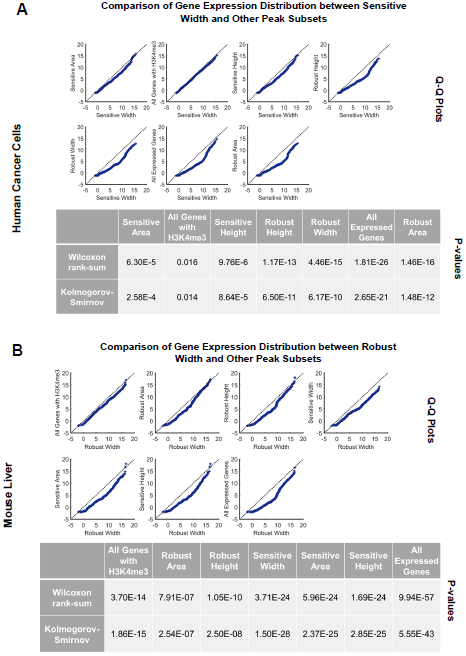
(Related to Fig 4) A. Q-Q plots, Wilcoxon rank-sum P-values and Kolmogorov-Smirnov P-values comparing distributions of gene expression levels associated with other peak subsets to the peak subset with sensitive width in human cancer cells. B. Same as in (A) but for robust width in mouse liver.

**Figure S14.**
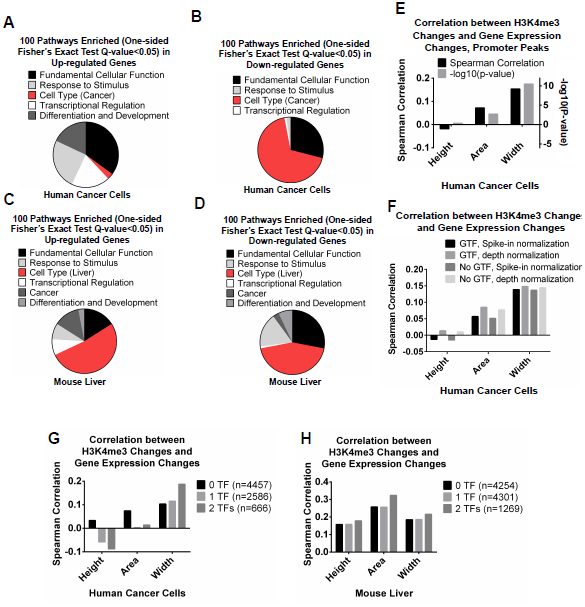
(Related to Fig 4) A. Annotation of top 100 pathways enriched in up-regulated genes in human cancer cells. B. Same as in (A) but for down-regulated genes. C. Annotation of top 100 pathways enriched in up-regulated genes in mouse liver. D. Same as in (C) but for down-regulated genes. E. Spearman correlation between H3K4me3 changes and gene expression changes with restriction of the analysis to promoter peaks in human cancer cells. F. Spearman correlation between H3K4me3 changes and gene expression changes using different methods for data analysis in human cancer cells. G. Spearman correlation between H3K4me3 changes and gene expression changes in genes associated with H3K4me3 peaks bound by 0, 1, or 2 TFs putatively associated with H3K4me3 width in human cancer cells. H. Same as (G) but for mouse liver.

**Figure S15.**
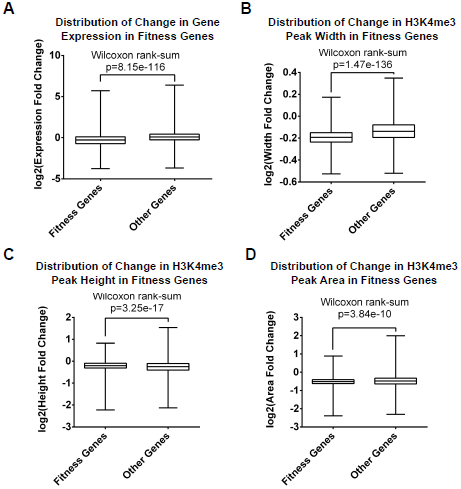
(Related to Fig 4) A. Distribution of gene expression changes in fitness genes (i.e. genes found to be essential for cancer cell survival and proliferation in a CRISPR-based screen) and all other genes. B. Distribution of width changes in H3K4me3 peaks associated with fitness genes and all other genes. C. Same as in (B) but for height. D. Same as in (B) but for area.

**Table S1.** Peak height, area, width and gene expression levels in different conditions.

